# AutoGeneS: Automatic gene selection using multi-objective optimization for RNA-seq deconvolution

**DOI:** 10.1101/2020.02.21.940650

**Authors:** Hananeh Aliee, Fabian Theis

## Abstract

Tissues are complex systems of interacting cell types. Knowing cell-type proportions in a tissue is very important to identify which cells or cell types are targeted by a disease or perturbation. When measuring such responses using RNA-seq, bulk RNA-seq masks cellular heterogeneity. Hence, several computational methods have been proposed to infer cell-type proportions from bulk RNA samples. Their performance with noisy reference profiles highly depends on the set of genes undergoing deconvolution. These genes are often selected based on prior knowledge or a single-criterion test that might not be useful to dissect closely correlated cell types. In this work, we introduce *AutoGeneS*, a tool that automatically extracts informative genes and reveals the cellular heterogeneity of bulk RNA samples. AutoGeneS requires no prior knowledge about marker genes and selects genes by simultaneously optimizing multiple criteria: minimizing the correlation and maximizing the distance between cell types. It can be applied to reference profiles from various sources like single-cell experiments or sorted cell populations. Results from human samples of peripheral blood illustrate that AutoGeneS outperforms other methods. Our results also highlight the impact of our approach on analyzing bulk RNA samples with noisy single-cell reference profiles and closely correlated cell types. Ground truth cell proportions analyzed by flow cytometry confirmed the accuracy of the predictions of AutoGeneS in identifying cell-type proportions. AutoGeneS is available for use via a standalone Python package (https://github.com/theislab/AutoGeneS).

## Background

Bulk RNA samples are routinely collected and profiled for clinical purposes and biological research to study gene expression patterns in various conditions such as disease states. Such samples reflect mean gene expression across thousands of cells and thus mask cellular heterogeneity in a complex tissue. However, the knowledge of cell-type composition and their fractions helps to characterize molecular changes in diseased tissues that are important for the identification of disease-related cell types as well as the development of targeted drugs and therapies [21]. Studying the variation of cell-type composition also opens new avenues in analyzing a tremendous yet underexplored quantity of biomedical data that has already been collected in clinics. Therefore, a number of bulk deconvolution techniques have been proposed in the literature for analyzing cellular composition from mixture samples [12, 13, 16, 20, 28, 27, 34]. These techniques mainly rely on a so-called signature matrix consisting of the gene signatures chiefly of well defined cell types [2, 20]. Some of these techniques are tissue-specific and offer a list of marker genes that can be employed for the deconvolution of those tissues [1, 21, 23, 26, 29]. However, due to the advances in single-cell RNA-sequencing (scRNA-seq), many cell subtypes are still being discovered whose specific markers are yet unknown. Therefore, automatic techniques are greatly desired to specify marker genes. To fill this need, previous studies have explored general feature selection techniques that can be extended to almost any tissue type [12, 17, 25, 32]. These techniques typically rank genes in each population (or cell type) based on a single-criterion comparison (often q-value from a t-test) and select the top-ranked genes that differentiate one population from the others. However, when complex tissues contain highly similar cell types and, more specifically, when the reference profiles are noisy (as is often the case for scRNA-seq data [5]), the informative genes must be filtered based on additional criteria. Moreover, these techniques often perform feature selection on each population individually and accumulate population-specific features. This can result in a large, overlapping set of genes between closely related populations. Therefore, it is important to globally select a set of markers that differentiate all populations.

In this study, we present a novel method, named AutoGeneS, that automatically selects genes for bulk deconvolution and accurately infers cell-type abundance. AutoGeneS is robust against technical and biological noise and can reveal cell abundance of complex mixture samples without any a priori knowledge about the marker genes. To accomplish this, we developed a feature selection method employing multi-objective optimization that leverages a signature matrix from single-cell or bulk-sorted transcriptional data. The optimization approach targets closely related cell types and aims at selecting a set of genes that simultaneously minimizes the correlation and maximizes the distance between the cell types. Through comprehensive benchmark evaluations and applications of expression in human samples of peripheral blood mononuclear cells (PBMC) with single-cell and bulk-sorted reference profiles, we show that selecting genes for deconvolution using AutoGeneS outperforms other approaches.

## Results

### AutoGeneS enables tissue characterization using reference profiles across multiple technologies

To dissect cellular content from bulk RNA samples, we first required reference gene expression profiles (GEPs), which may originate from various technologies such as single-cell experiments or sorted cell populations (Figure 1**a**). To derive reference GEPs from scRNA-seq data, dimensionality reduction and clustering were first performed to reveal cell types and annotate cells. The GEP of a cell type was then selected as the closest point to the centroid of the corresponding cluster (see Methods for technical details). AutoGeneS is a method that takes GEPs and identifies informative genes for deconvolution using multi-objective optimization (see next section). This was particularly crucial for (i) reducing computational complexity by eliminating unexpressed genes or genes with a uniform expression level across the cell types that is important for large scale studies and (ii) increasing the signal-to-noise ratio by selecting informative genes that differentiate the cell types. Once we filtered the reference GEPs for informative genes, called a *signature matrix*, we leveraged them to predict cell proportions from other mixture profiles.

**Figure 1:**
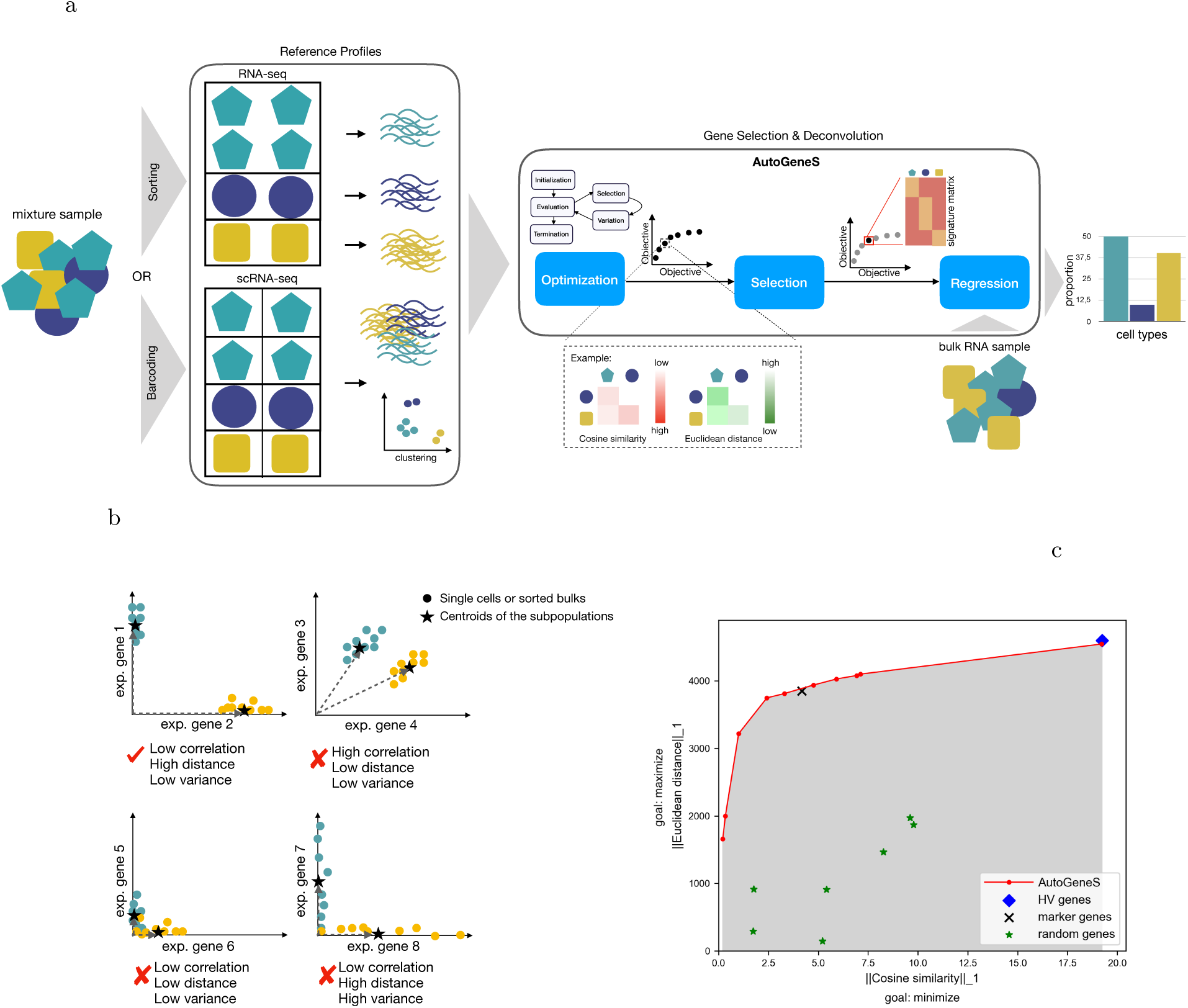
Framework for dissecting cellular composition. **a** The workflow starts by transcriptome profiling of single cells or sorted cell populations to generate reference profiles. For scRNA-seq, cells are clustered, and the mean expressions of populations are computed as reference profiles. AutoGeneS is then applied on the reference profiles to find genes that discriminate cell populations and measure proportions. AutoGeneS is a multi-objective optimization-based approach using evolutionary algorithms (discussed in Methods). During optimization, several solutions, each consisting of a set of genes, are generated and evaluated for multiple objectives (e. g., correlation and distance). Finally, the solutions that cannot be improved for any of the objectives without degrading at least one of the other objectives are returned. Without additional subjective preference, all of these solutions are considered equally good, and one is selected for regression (for discussion, see Methods). The reference profiles are then filtered for the genes in the selected solution called the signature matrix. The cell-type proportions of a bulk RNA sample are then inferred using a regression technique like Nu-SVR. **b** Finding discriminating genes (also known as feature selection) is one of the main challenges in bulk deconvolution. Here, we indicate through four examples that minimizing inter-population correlation, the criterion which has been employed most frequently in the literature, is not necessarily sufficient. In the first example, gene 1 and gene 2 are ideal for deconvolution because they are expressed in only one of the two populations, and the populations have a high distance and a low intra-population variance. However, gene 3 and gene 4 may not be selected because the populations are correlated in those dimensions. Moreover, gene 5 and gene 6 are not appropriate because the centroids of the populations are very close, and the proportion estimates become very sensitive to smaller changes due to possible noise in the data. Similarly, gene 7 and gene 8 are not good candidates because of the high variance in both populations. In this last case, the centroids are not representative of the cell populations. **c** The plot illustrates the objective values for Pareto-optimal solutions using AutoGeneS for human embryonic data from [8] and compares these solutions with those using highly variable (HV) genes, marker genes, or random sets of genes. The AutoGeneS solutions dominate all the other solutions.

Previous studies show that the expression profiles from each cell type are linearly additive; this makes the contribution of each cell type proportional to its fraction in the mixture profile [34]. Hence, we then employed Nu-Support Vector Regression (Nu-SVR) to deconvolve a given mixture as the last step of the workflow. This makes AutoGeneS robust against technical and biological noise by discarding a certain fraction of outliers specified by *Nu* and performing regression on the remaining samples [7] (discussed in Methods).

### Multi-objective optimization can learn non-collinear genes

In this study, we propose a multi-objective optimization approach to select a set of non-collinear genes and show that excluding collinear genes significantly improves the performance of deconvolution approaches. Collinearity (also called multicollinearity) occurs when two or more predictor variables (cell-type-specific GEPs) in a statistical model (usually in a regression-type analysis) are linearly correlated. High collinearity means that collinear predictors share a substantial amount of information, which can cause model non-identifiability. In this case, coefficient estimates (the proportions of cell types) can be distorted by inflated standard errors that swing widely and are highly sensitive to small changes in the dataset. Because biological datasets (specifically scRNA-seq data) are typically considered noisy, collinearity is one of the main challenges in bulk deconvolution. Several quantifications of collinearity are proposed in the literature, with the most common being the pair-wise correlation coefficient [11]. While correlation and collinearity are not equivalent, high absolute correlation coefficients are usually indicative of high linear relatedness. For example, when the correlation between two predictors is zero, they are orthogonal, and their collinearity is zero as well. Auto-GeneS is then developed to select a set of genes that minimizes the global correlation between all the predictors simultaneously to reduce collinearity. By global correlation, we mean the sum of the pair-wise correlation coefficients between all pairs of populations.

However, selecting a subset of genes that only minimizes the correlation might not be enough (Figure 1**b**). Indeed, when the Euclidean distance between the centroids of two populations is low, the coefficient estimates become highly sensitive to noise in the dataset (Figure 1**b**, genes *g*_5_ and *g*_6_). Moreover, selecting unstable genes with high inter-cluster variance can result in low representability of the centroids, which can increase regression error (Figure 1**b**, genes *g*_7_ and *g*_8_). Therefore, rather than selecting genes based only on a pair-wise correlation coefficient, we propose a multi-objective optimization approach that simultaneously minimizes global correlation and maximizes the sum of pair-wise Euclidean distances (Figure 1**a**, AutoGeneS framework). We may also consider the inter-cluster variance as a third objective to be minimized, but instead we recommend adding this as a constraint to the optimizer by filtering out genes with high inter-cluster variance (for more discussion, see Methods).

For a multi-objective optimization problem, there usually exists no single solution that simultaneously optimizes all objectives. In this case, the objective functions are said to be conflicting, and there exists a (possibly infinite) number of Pareto-optimal solutions. Pareto-(semi)optimal solutions are a set of all solutions that are not dominated by any other explored solution. Figure 1**c** illustrates a representative set of non-dominated solutions using AutoGeneS for human embryonic scRNA-seq data from [8] (discussed later in this section) on 4,000 preselected HV genes. Each solution in this figure refers to a set of genes that forms a signature matrix. Conventional deconvolution techniques typically use a t-test between each population and all other populations to obtain differentially expressed genes called *marker genes*. Figure 1**c** compares the objective values for AutoGeneS, marker genes, HV genes, and a set of random solutions, for which some genes are randomly selected as markers. The plot shows that the solutions from AutoGeneS dominate other solutions and, more importantly, that AutoGeneS is capable of finding solutions with a very low global correlation and a relatively high global distance. Pareto-optimal solutions offer a set of equally good solutions from which to select, depending on the dataset (discussed in Methods).

### Evaluation on simulated bulk tissues

First, to systematically benchmark, we applied AutoGeneS to synthetic bulk RNA-seq data. The reference data for signature learning was human embryonic scRNA-seq data from [8]. To make the deconvolution challenging but also more realistic, the synthetic bulk RNA samples were generated by summing the sorted bulk RNA-seq read counts of the same tissue from [8]. Indeed, library preparation protocols vary across different sequencing technologies as well as laboratories, and bulk deconvolution analysis therefore may be performed on reference data generated differently in either way. In this scenario, true cell-type proportions are known, which allows a comparison of the accuracy of the proposed technique. In this experiment, AutoGeneS searched for 400 marker genes that minimized cosine similarity (as a measure of correlation) and maximized Euclidean distance between six major cell types (Figure 2). After filtering the 400 marker genes with the lowest mean correlation coefficients among all the Pareto-optimal solutions, the average pair-wise correlation coefficient was equal to 0.07. This shows that AutoGeneS efficiently explores the search space seeking non-collinear genes (Figure 2**b**). Moreover, compared with ground truth cell proportions, the deconvolution results show high accuracy, with a p-value < 1e-78 (Figure 2**c**). The heatmap of the signature matrix further indicates that the optimization result converges to the genes that are either highly expressed in only one cell type or lowly but differentially expressed between the cell types (Figure 2**d**). More details on synthetic bulk generation are explained in the Methods section.

**Figure 2:**
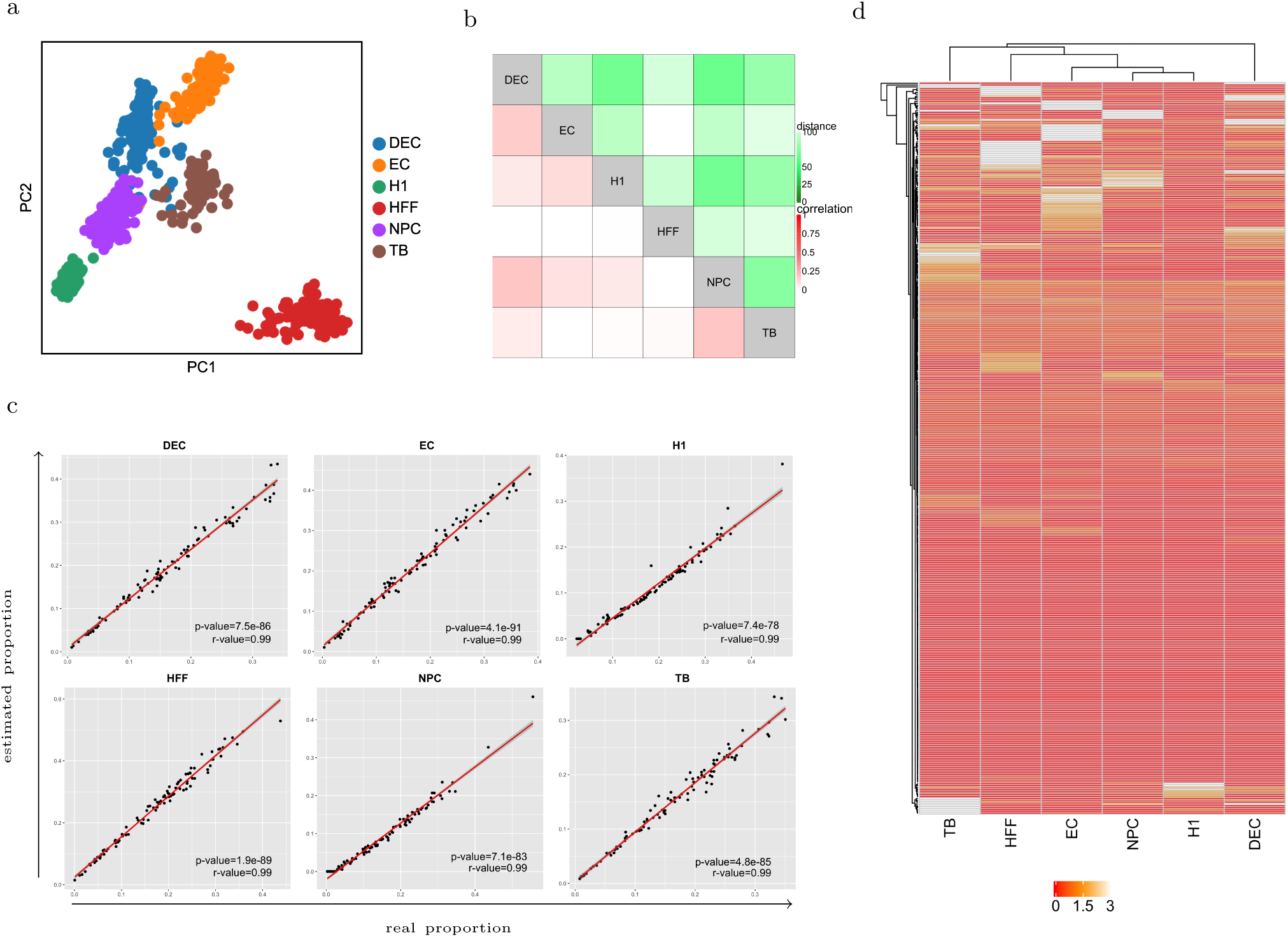
Bulk Deconvolution of synthetic bulk RNA samples with a human embryonic single-cell reference profile. Synthetic samples were generated by summing the sorted bulk RNA-seq read counts of the same tissue. **a** Principal component analysis (PCA) projection of the scRNA-seq data with six major populations. Shown is PC1 vs. PC2. **b** Correlation and distance matrices of the signature matrix after optimization. The correlation between the cell types after optimization is negligible (average pair-wise correlation coefficient of 0.07). **c** Scatter plots illustrating real and estimated proportions for 100 synthetic bulk RNA samples. The significance of the results were assessed by r-values and p-values from linear regression (solid red lines). Linear regression shows highly significant correlation between estimated and real proportions. **d** Heat map of signature matrix using AutoGeneS for gene selection. AutoGeneS converges to the genes that are highly expressed only in one cell type and lowly but differentially expressed between cell types. NPC, neuronal progenitor cell; DEC, definitive endoderm cell; EC, endothelial cell; TB, trophoblast-like cell; HFF, human foreskin fibroblasts.

### Deconvolution of PBMC samples with sorted bulk RNA-seq

We next deconvolved bulk RNA-seq data of PBMC samples from [23]. The bulk RNA-seq data consisted of gene expression measurements of 12 individuals (details in Methods). The sorted bulk RNA samples of 29 immune cell types in the blood samples of four individuals were also available from [23] to build the signature matrix. The flow cytometry proportions of each sample were employed for validating the deconvolution results. The authors in [23] delineated the cell types with the highest mean Pearson correlation and merged those from the classification used for FACS (i. e., the 29 cell types) with no detectable and/or no specific signal into broader cell types. From this procedure, they selected 17 cell types that we also used in our analysis. Besides the sorted bulk RNA samples, the authors in [23] used as reference the mixture samples weighted by flow cytometry proportions. Because signatures are predicted from the same samples being used for deconvolution, we referred to it as a supervised approach that other unsupervised approaches might hardly exceed. In this analysis, we compared four unsupervised feature selection methods: AutoGeneS, HV genes, marker genes based on *t-test* (T-MG), and marker genes based on condition number (C-MG) with t-test-based preselection (as used in CIBERSORT [25] and Shiny [23]). AutoGeneS successfully found the markers that minimize the correlation while simultaneously maximizing the distance between each pair of cell types (Figure 3**a**). The linear agreement between the real proportions from flow cytometry and the predicted ones was determined by correlation coefficient (r-value) and two-sided p-value (the direct comparison of AutoGeneS and flow cytometry results is presented in Figure S2). As shown in Figure 3**b**, existing unsupervised methods failed to capture the correct proportions for several cell types; therefore, AutoGeneS outperforms them. Moreover, although the results of AutoGeneS are close to those of supervised Shiny, AutoGeneS used only 500 genes, while Shiny used 1296 (Figure 3**c**). Using fewer genes significantly reduces the computational time of the analysis (Figure 3**c**, Table 1).

**Table 1:**
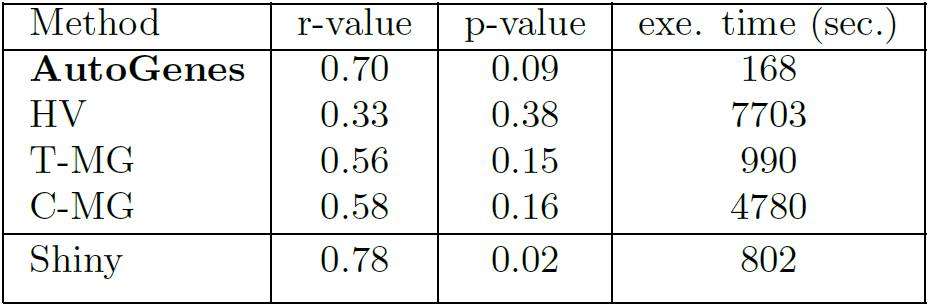
Summary of performance

**Figure 3:**
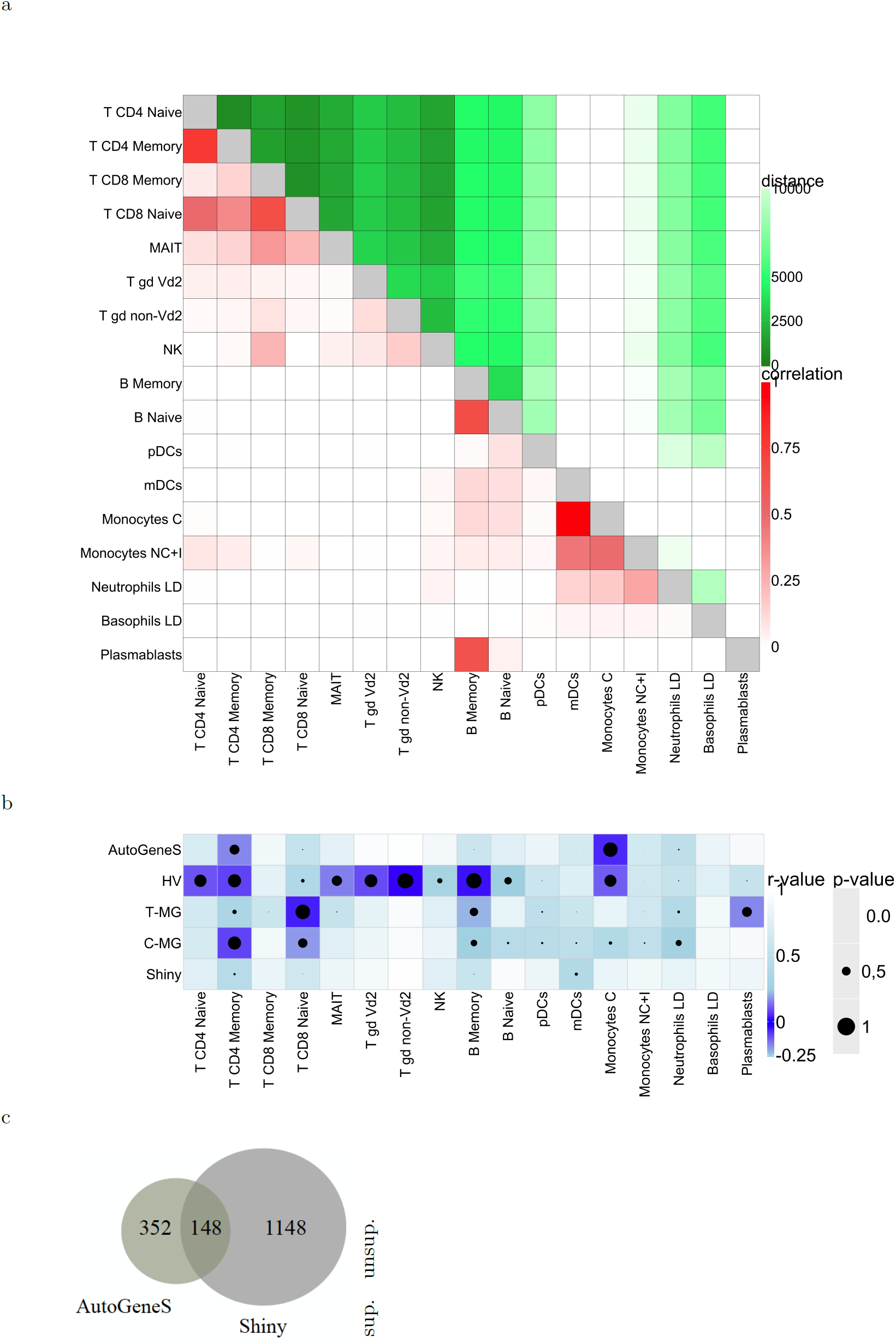
Bulk Deconvolution of 12 PBMC samples with bulk-sorted reference profiles. **a** Correlation and distance matrices of the signature matrix after optimization. With the exception of naturally closely correlated cell types (like T cells and monocytes), the correlation between the cell types is negligible (average pair-wise correlation coefficient = 0.07). **b** p-values and r-values for various feature selection techniques indicate the efficiency of AutoGeneS compared to existing works. For each technique and cell type, the p-values and r-values were measured by fitting a linear regression with estimated and real proportions from flow cytometry (the mean r-value and p-value are represented in Table 1). The regression results using AutoGeneS are represented in Supplementary Figure S2. **c** The number of joint features among AutoGeneS and Shiny. Shiny, a supervised feature selection technique, used 1296 genes and AutoGeneS was set to select 500 genes in an automatic manner for deconvolution. This decrease in genes improved the computational time of AutoGeneS by up to 46 times. The execution times are summarized in Table 1.

### Deconvolution of PBMC samples with scRNA-seq

We next turned to the problem of inferring cell-type-specific GEPs when bulk and single-cell reference data originate from different studies. We deconvolved PBMC samples from [23] using the scRNA-seq data of blood from [3]. The single-cell data consists of cell types including: adipocytes, B cells, CD4+ T cells, CD8+ T cells, chondrocytes, dendritic cells, endothelial cells, eosinophils, erythrocytes, hematopoietic stem cells, macrophages, monocytes, natural killer (NK) cells, neutrophils, and skeletal muscle. We filtered out populations with fewer than 45 cells and were left with six major cell types. NK cells, CD4+ T cells, and CD8+ T cells have a high correlation that is also visible in the uniform manifold approximation and projection (UMAP) [22] of the single-cell data (Figure 4**a**). Here, we observed how the aforementioned unsupervised feature selection techniques dealt with high correlation. Compared with the ground truth cell proportions as determined by flow cytometry, the low correlation in the deconvolution results might be driven either by platform-specific variation between bulk and scRNA-seq data or the inability of the feature selection techniques to extract informative genes. To validate the former hypothesis, we deconvolved the bulk RNA samples using single-cell reference data with the list of genes selected in the supervised manner using Shiny. The existing unsupervised methods failed to infer the proportion of neutrophils and CD4+ T cells, and AutoGeneS outperformed them. Also, AutoGeneS had the closest results to Shiny while even predicting the fraction of NK cells more accurately (Figure 4**c**). Direct comparison of AutoGeneS and flow cytometry results is presented in Figure S3.

**Figure 4:**
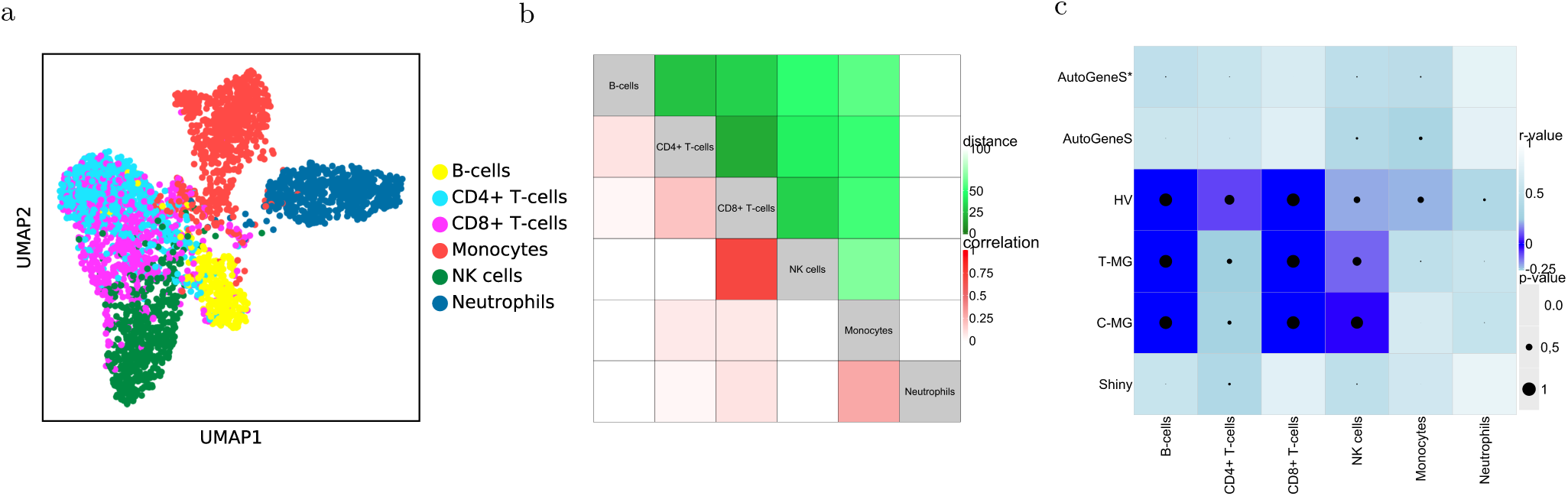
Bulk Deconvolution of 12 PBMC samples with a single-cell reference profile. **a** UMAP of PBMC scRNA-seq data with six major cell types shows high correlation between CD4+ T cells, CD8+ T cells and Natural Killer (NK) cells. **b** Correlation and distance matrices of the signature matrix after optimization. Except for CD8+ T cells and NKs, the correlation between the cell types is negligible (average pair-wise correlation coefficient = 0.10). Moreover, because CD4+ and CD8+ T cells are naturally close to each other, the distance between them is also small. **c** p-values and r-values for various feature selection techniques indicate that basic AutoGeneS and hierarchical AutoGeneS (AutoGeneS*) outperform other unsupervised methods. The p-values and r-values were measured by fitting a linear regression with estimated and real proportions from flow cytometry. The regression results using hierarchical AutoGeneS is represented in Supplementary Figure S3.

### Hierarchical optimization for highly correlated cell types

The correlation matrix reflects the similarity between cell types. Based on this, the user then groups the cells with almost no differentiating signals. After running the optimization, the correlation matrix on the selected features reveals whether some of the cell types are still correlated. If the correlation is high for some cell types, we propose running AutoGeneS at a different stage on those correlated cell types and aggregating the new features with the earlier ones to build the final signature matrix. For the dataset studied in Figure 4, we ran AutoGeneS separately for CD4+ and CD8+ T cells and added in total 10 more genes to the signature matrix that can better differentiate between these two cell types (adding more genes can increase the correlation between other cell types). The results show that the hierarchical AutoGeneS (AutoGeneS*) obtains better results compared with basic AutoGeneS specifically for CD8+ T cells and monocytes (Figure 4**c**).

## Discussion and Conclusions

Knowledge of cell-type compositions is important to reveal cellular heterogeneity in diseased tissues and is helpful to identify the targets of a disease. Although Bulk RNA-seq averages gene expression across thousands of cells and thus masking cellular heterogeneity in complex tissues, scRNA-seq usually does not reflect true cell-type proportions in intact tissues. Furthermore, it remains costly for use in clinical studies with large cohorts. Therefore, several bulk deconvolution techniques have been proposed in the literature to predict cell-type proportions from mixed samples. Existing approaches either rely on preselected marker genes or perform a single-criterion test to identify differentially expressed genes among cell types. In this study, we introduce AutoGeneS which, for the first time, selects informative genes based on multiple criteria and infers accurate cell-type proportions. AutoGeneS selects informative genes using a multi-objective optimization approach that simultaneously minimizes average pair-wise correlation coefficients and maximizes the Euclidean distance between cell types. By minimizing correlation, we reduce the collinearity as one of the main challenges in bulk deconvolution. We simultaneously ensure that the Euclidean distance between the cell types is maximized to safeguard predictions against noise. Through comprehensive benchmark evaluations and analysis of multiple real datasets, we show that AutoGeneS outperforms other approaches. Moreover, ground truth cell-type proportions identified by flow cytometry confirm the accuracy of the predictions by AutoGeneS.

While in this study we use correlation and distance as objectives, AutoGeneS offers a flexible framework that can be easily extended to other user-specific objectives. Among the unsupervised feature selection techniques previously proposed for bulk deconvolution, we implemented two objective functions inspired by the works in [12, 24]. (i) Newman et al [24] proposed first preselecting top G marker genes for each population using a t-test with a low q-value and later iterating G across all populations and retraining the signature matrix with the lowest condition number. We studied a new implementation of this approach by integrating condition number into AutoGeneS as an objective function applied on either HV genes or preselected marker genes. (ii) MuSic [12] is another approach that weights genes according to their variance before running deconvolution on a whole gene set. For single-cell experiments, we can consider the sum of the variances of populations as an objective to be minimized. However, selecting genes based on these objectives, we did not notice an improvement in the results (data not shown).

AutoGeneS currently requires reference profiles of desired cell types to search for informative genes. The reference profiles can be provided by either sorted bulk RNA-seq or scRNA-seq. Using scRNA-seq data has the advantage of putatively revealing novel cell types and subtypes in a system that no prior knowledge of marker genes for such novel populations exists. We have demonstrated in this work that AutoGeneS can successfully utilize scRNA-seq data as effective references to identify marker genes for differentiating cell types.

## Methods

### Simulated bulk RNA sample generation

To validate the proposed deconvolution process, we required bulk RNA-seq with known cell-type proportions. In total, 100 synthetic bulk profiles were generated as a mixture of reference-sorted bulk RNA samples with known fractions. Because the reference profiles and the synthetic bulk RNA samples were generated using different sequencing technologies, this was a more realistic way of generating synthetic bulk RNA samples compared with the approaches that subsample scRNA-seq data and sum the expression profiles to simulate bulk RNA samples.

### Deconvolution

To infer cell-type abundance, related work generally assumes that the count matrix *Y* of *m* mixture samples with *N* genes is the weighted sum of *K* cell-type-specific count profiles with the same *N* genes, represented by matrix *X* [21, 13, 15]. This can be modeled as a system of linear equations as follows:

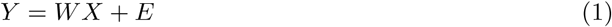

where, *W* is an *m* × *k* fractional abundance matrix and *E* represents the residual errors. To compute *W*, we solve the equation for only a subset of genes *g* ≪ *N* selected using AutoGeneS:

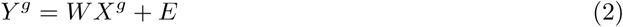

To build the signature matrix using bulk-sorted RNA-seq, we averaged transcripts-per-million (TPM)-normalized cell-type-specific profiles in non-log linear space and filtered the markers. For scRNA-seq, we tested different techniques to define the centroid of a cell type such as the median or the mean of all samples, the mean of samples in the densest area of a cluster, or the mean of subsamples. However, with the exception of the median of a group with poor performance, we observed no significant difference in deconvolution performance when applying different techniques (data not shown). Therefore, we used the mean expression of all cells in a population after normalizing each cell into TPM and filtered the markers to obtain population-specific reference signals.

### Multi-objective Optimization

The proposed feature selection approach solves a multi-objective optimization problem that can be generally formulated as:

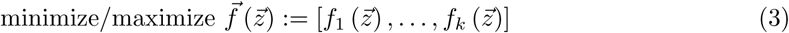

subject to:

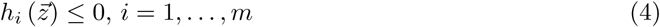

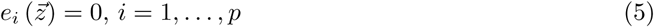

where 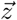 is the vector of decision variables and *f*_*i*_ : ℝ^*n*^ → ℝ, *i* = 1, …, *k* are the objective functions which evaluate the quality of a solution by assigning a fitness value to it. Also, *h*_*i*_, *e*_*j*_ : ℝ^*n*^ → ℝ, *i* = 1, …, *m, j* = 1, …, *p* are referred to as the inequality and equality constraint functions of a problem which must be satisfied. In AutoGeneS, we have *n* binary decision variables where *n* is equal to the number of genes from which the optimizer selects the markers. The value of a decision variable represents whether the corresponding gene is selected as a marker. Later, we evaluate the objective functions (correlation and distance) only for genes *G* whose decision variables are set to one. Considering 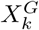 as the expression profile of cell type *k* for genes *G*, the correlation objective is measured as:

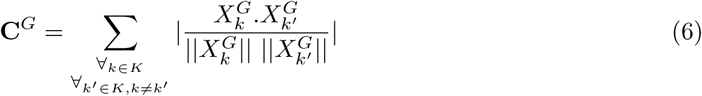

and the distance objective is measured as:

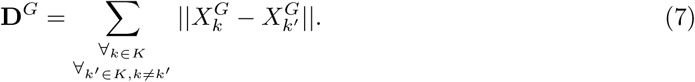

AutoGeneS allows setting the number of selected markers G as a fixed value that can be implemented as a constraint (all the solutions where |*G*| is larger than a desired value, are marked as infeasible and will not be evaluated). However, we show later how this constraint is implemented differently in AutoGeneS to obtain a better performance.

Several techniques have been proposed in the literature for solving multi-objective optimization problems. Among these, meta-heuristics such as multi-objective evolutionary algorithms (MOEAs) are quite popular and well established mainly because of their flexibility and widespread applications [10, 9, 30]. MOEA denotes a class of search methods where the decisions are made in the presence of trade-offs between objectives. As the name suggests, multi-objective optimization involves more than one objective function to be optimized at once. When no single solution exists that simultaneously optimizes each objective, the objective functions are said to be conflicting. In this case, the optimal solution of one objective function is different from that of the others. This gives rise to a set of trade-off optimal solutions popularly known as *Pareto-optimal* solutions. The list of Pareto-optimal solutions includes non-dominated solutions, which are explored so far by the search algorithm. These solutions cannot be improved for any of the objectives without degrading at least one of the other objectives. Without additional subjective preference, all Pareto-optimal solutions are considered to be equally good.

AutoGeneS uses a genetic algorithm (GA) as one of the main representatives of the family of MOEAs. GA uses a population-based approach where candidate solutions that represent individuals of a population are iteratively modified using heuristic rules to increase their fitness (i. e., objective function values) [14, 19]. The main steps of the generic algorithm are as follows:

1. Initialization step: Here the initial population of individuals is randomly generated. Each individual represents a candidate solution that, in the feature selection problem, is a set of marker genes. The solution is represented as a bit string with each bit representing a gene. If a bit is one, the corresponding gene is selected as a marker.
2. Evaluation and selection step: Here the individuals are evaluated for fitness (objective) values, and they are ranked from best to worst based on their fitness values. After evaluation, the best feasible individuals are then stored in an archive according to their objective values.
3. Termination step: Here, if the termination conditions (e. g., if the simulation has run a certain number of generations) are satisfied, then the simulation exits with the current solutions in the archive. Otherwise, a new generation is created.
4. If the simulation continues, the next step is creating offspring (new individuals): The general GA modifies solutions in the archive and creates offspring through random-based *crossover* and *mutation* operators. First, parents are selected among the candidates in the archive. Second, the crossover operator combines the bits of the parents to create the offspring. Third, the mutation operator makes random changes to the offspring. Offspring are then added to the population, and the GA continues with step 2.

The algorithms in Figure S1 represent how the crossover and mutation operators are implemented in AutoGeneS to always return individuals with a fixed number of marker genes. In our feature selection problem, we are generally interested in finding the most informative genes out of *N* variable genes (e. g., 4,000 HV genes). This results in a search space with 2^|*N*|^ possible solutions. However, the number of possible solutions is decreased to 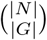 when we only search for solutions with a fixed number of genes. Therefore, this implementation avoids generating infeasible solutions (where the number of markers are not equal to |*G*|) and significantly improves the efficiency of the whole GA. We recommend to select |*G*| ∈ [300 − 500] genes depending on the number of cell types. For the problems in Figures 2 and 3, we searched for 400 marker genes and for the problem in Figure 4, 300 marker genes out of 4,000 HV genes were selected.

### Signature gene selection

By their nature, multi-objective optimization problems give rise to a set of Pareto-optimal solutions that need further processing to find a single solution that satisfies the subjective preferences of the users. AutoGeneS also plots the set of Pareto-optimal solutions; comparing the correlation and the distance of the two extremes (the solutions with lowest correlation and highest distance) indicates which solution might be better. In our analysis, we picked the solution with the lowest correlation.

### Highly variable genes

Highly variable (HV) genes are those that are informative of the variability in a dataset [5]. These genes are usually filtered as the first step of feature selection. Depending on the complexity of a dataset, typically between 1,000 and 5,000 HV genes are selected for downstream analysis. The authors in [18] suggest that downstream analysis is only as robust as the exact choice of the number of HV genes. Due to the possible batch effects especially regarding varying library preparation protocols between reference and mixture samples, we prefer to select a higher number of HVGs, like 4,000. We used the method implemented in single-cell analysis in Python (SCANPY) [33] for selecting HV genes in which genes are binned by their mean expression and those with the highest variance-to-mean ratio are selected as HV genes in each bin. Of course, before applying this method, genes with a low average expression are excluded (if not already done for the external datasets) as a quality control filter.

### Marker genes

To find differentially expressed or marker genes in a population of interest, statistical tests like Wilcoxon rank-sum test or t-test are usually used to rank genes by their difference in expression between two groups (the population of interest and the remaining populations). In the present study, for both T-MG and C-MG, we have used t-test from the Seurat package [6, 31] with Benjamini-Hochberg (BH) corrected p-values [4] (called q-values) on log-transformed data. The top G marker genes with lowest q-values are selected for each population and aggregated as T-MG. We have selected G so that T-MG contained in total about 500 genes. Selecting C-MG has two more steps: (i) sorting the top G genes by decreasing fold change compared to other cells and (ii) and iterating G from 50 to 200 across all populations and retaining the signature matrix with the lowest condition number (as proposed in [24]).

### Regression

Although the feature selection strategy is general and orthogonal to regression techniques, we used Nu-SVR for deconvolution. Compared with regular linear regression, which attempts to minimize regression error, support vector regression (SVR) tries to fit the regression error within a certain threshold. The *ϵ*-loss function in SVR ignores errors that are within *ϵ* distance of the observed value by treating them as equal to zero. For other points outside this boundary, the loss is measured based on the distance between observed value *y* and the *ϵ* boundary. This produces a better-fitting model by highlighting the loss of outliers alone. In Nu-SVR, Nu ∈ (0, 1] indicates an upper bound on the fraction of training errors (poorly predicted samples) and a lower bound on the fraction of support vectors to use. The regression is performed on normalized, non-log-transformed data. After the regression, we ensure that the weights are positive and they sum to one. Thus, we set the negative values to 0 and normalize the rest to sum to one.

The execution times underlying Figures 3**c** were measured on an Intel(R) Xeon(R) Gold 6126 CPU at 2.60GHz with 395GB of RAM.

### Normalization

The external RNA-seq datasets were downloaded and analyzed using the authors’ normalization consisting of TPM or reads per kilobase of transcript per million (RPKM). The count matrix for scRNA-seq datasets were normalized to one million counts per cell.

## Data Availability

In this study, we have used existing datasets which are openly available as cited in the References section. The source data underlying Figure 2 are downloaded from GEO with accession code GSE75748. The 12 bulk RNA samples as well as sorted bulk samples underlying Figures 3, 4 and S2 are downloaded from GEO with accession codes GSE106898 and GSE10701, respectively. Finally, the scRNA-seq blood data underlying Figures 4 and S3 is accessible via [3].

## Code Availability

AutoGeneS is publicly available as a Python package at Github: https://github.com/theislab/AutoGeneS. A tutorial and an example are also provided.

## Acknowledgment

We would like to thank the many people who proofread the case study notebook and the manuscript and improved it with their comments and expertise. For this, we acknowledge the input of Carsten Marr, Anna Wojtuszkiewicz, Benjamin Schubert, and Matthias Heinig. This work was supported by the Helmholtz Association, Incubator grant sparse2big.

## Supplementary Material

**Figure S1:**
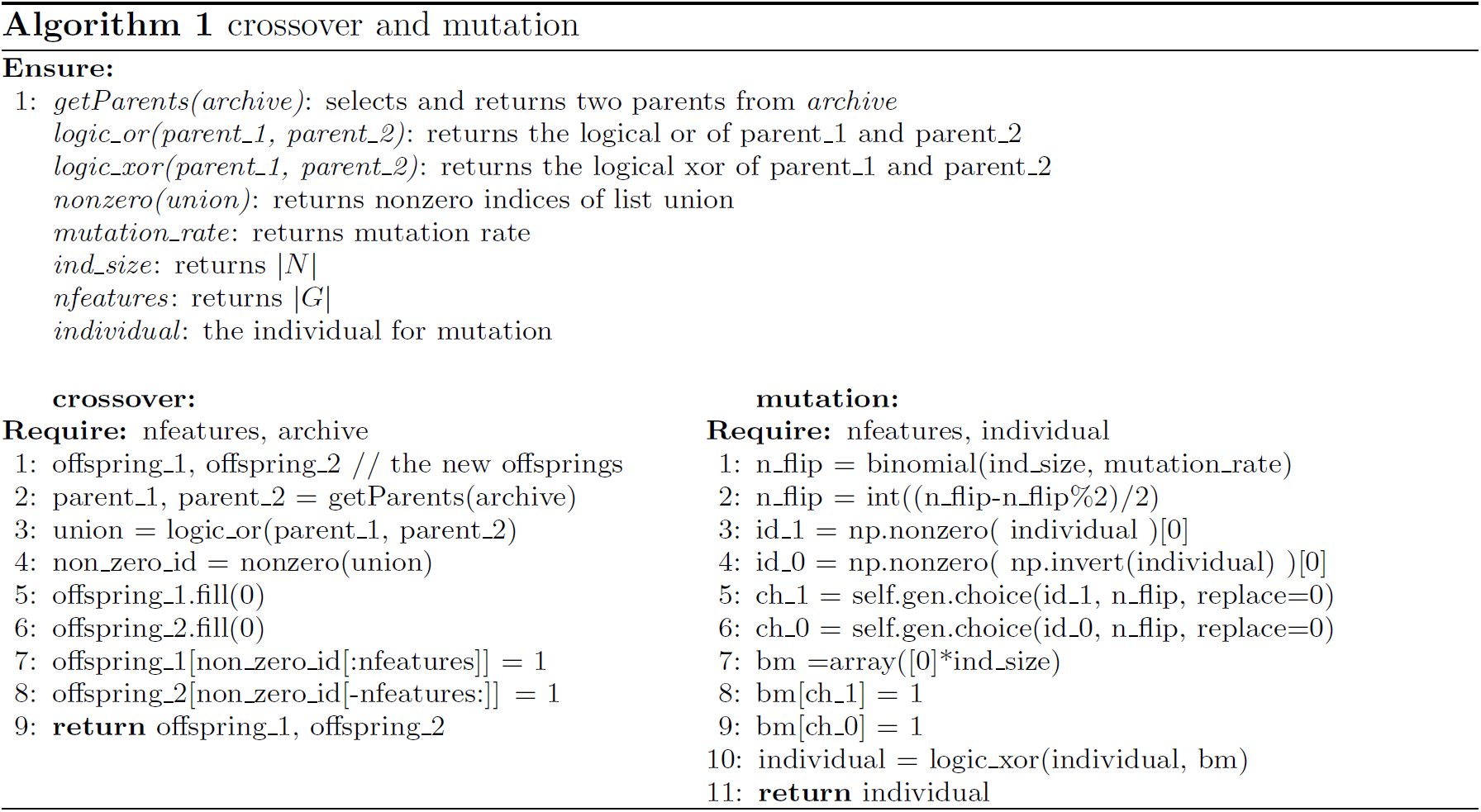
Crossover and mutation algorithms. The crossover operator combines the bits of the parents to generate new offspring with a fixed number of active bits (the bits with a value of 1). First, the union of the parents is calculated, and the first resulting |*G*| active bits of that are stored as *offspring_*1. The last |*G*| active bits are stored as *offspring_*2. The mutation operator makes random changes in the newly generated offsprings. It first selects the number of mutations (*n_flip*) and then randomly flips *n_flip/*2 bits with a value of 0 and *n_flip/*2 bits with a value of 1 in each offspring.

**Figure S2:**
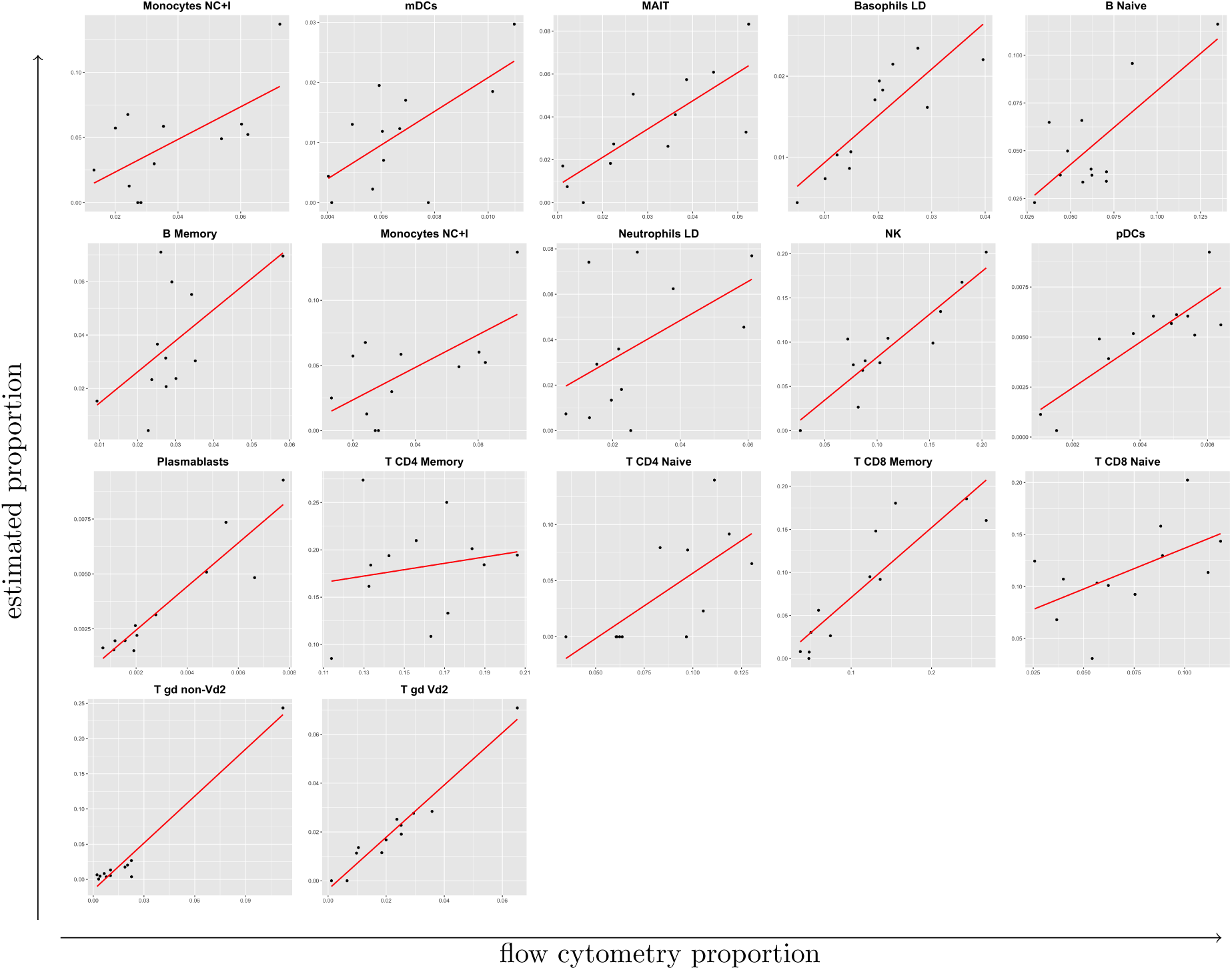
Deconvolution of individual cell types in 12 PBMC samples where sorted RNA-seq is used to generate the reference profiles. Shown is the direct comparison between AutoGeneS and flow cytometry for the indicated 17 cell types in PBMC samples. Concordance was determined by linear regression (solid red lines).

**Figure S3:**
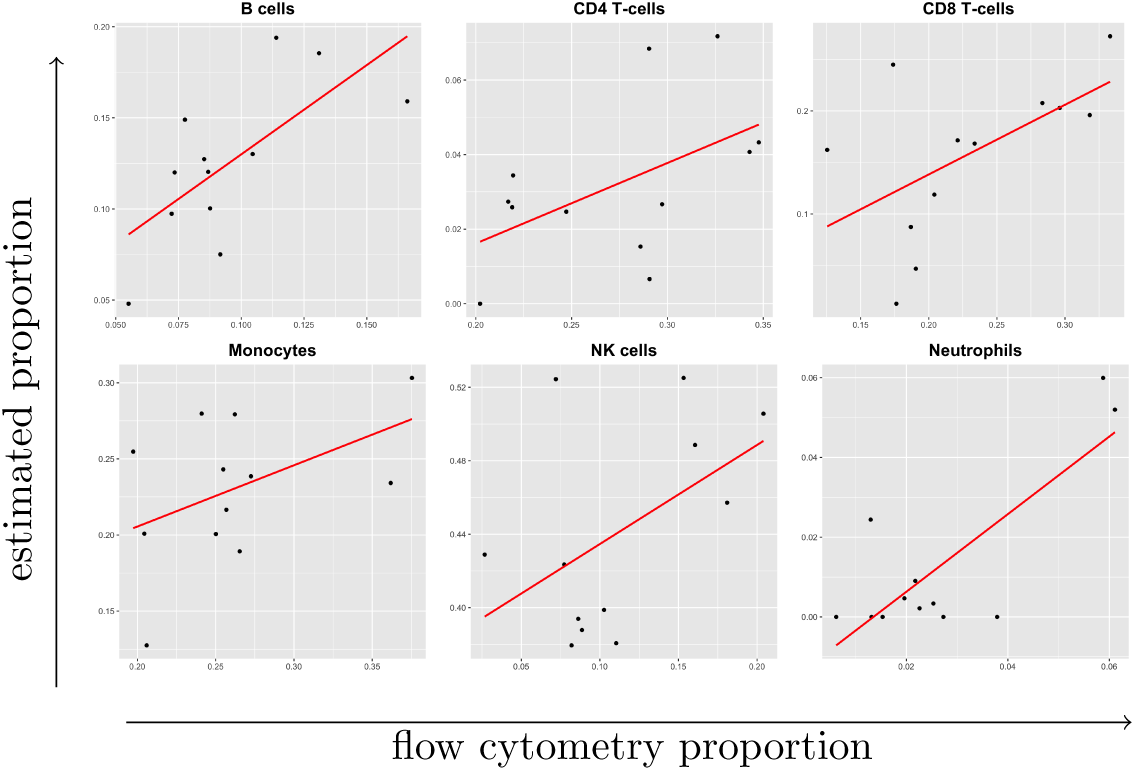
Deconvolution of individual cell types in 12 PBMC samples where single-cell RNA-seq is used to generate the reference profiles. Shown is the direct comparison between hierarchical AutoGeneS and flow cytometry for the indicated 6 major cell types in PBMC detected at single-cell resolution. Concordance was determined by linear regression (solid red lines).

